# Macroevolution of dimensionless life history metrics in tetrapods

**DOI:** 10.1101/520361

**Authors:** Cecina Babich Morrow, S. K. Morgan Ernest, Andrew J. Kerkhoff

**Affiliations:** Department of Biology, Kenyon College, Gambier, OH, USA; Center for Biodiversity and Conservation, American Museum of Natural History, New York, NY, USA; Department of Wildlife Ecology and Conservation, University of Florida, Gainesville, FL, USA

## Abstract

Life history traits represent organism’s strategies to navigate the fitness trade-offs between survival and reproduction. Eric Charnov developed three dimensionless metrics to quantify fundamental life history trade-offs. Lifetime reproductive effort (LRE), relative reproductive lifespan (RRL), and relative offspring size (ROS), together with body mass, can be used classify life history strategies across the four major classes of tetrapods: amphibians, reptiles, mammals, and birds. First, we investigate how the metrics have evolved in concert with body mass. In most cases, we find evidence for correlated evolution between body mass and the three metrics. Finally, we compare life history strategies across the four classes of tetrapods and find that LRE, RRL, and ROS delineate a space in which the major tetrapod clades occupy mostly unique subspaces. These distinct combinations of life history strategies provide us with a framework to understand the impact of major evolutionary transitions in energetics, physiology, and ecology.

## Introduction

Life history traits quantify the two crucial components of fitness: survival and reproduction. A species’ life history strategy, i.e. how it allocates resources to survival and reproduction, impacts its fitness and thus its success in terms of population growth and extinction risk (Sol et al. 2012; Capellini et al. 2015; Allen et al. 2017). Because resources are often limited, organisms cannot optimize both individual survival and reproductive investment, so allocation to one component of life history necessitates trade-offs in other areas (Stearns 1989; Charnov and Downhower 1995). Organisms navigate these constraints in a variety of ways to maximize overall fitness, creating an astonishing diversity of life history strategies across the tree of life. This diversity of life history strategies makes it challenging to assess the impacts of major evolutionary transitions: disparate evolutionary lineages may vary substantially in their reproductive physiologies and behaviors, schedules of growth and development, and pace and span of life. Comparing life histories across such varied lineages requires an approach that distills complex life histories into comparable and biologically meaningful metrics.

Life history theory is an attempt to understand how natural selection generates this diversity based on fundamental trade-offs in survival and reproduction. In a series of publications (Charnov and Berrigan 1991; Charnov 1993, 2002; Charnov and Downhower 1995), Eric Charnov developed a general model for life history evolution based on four basic premises: 1) the environment dictates the expected adult mortality rate for an organism; 2) natural selection acts on the age at first reproduction to maximize lifetime reproductive success given that mortality rate; 3) once organisms reach reproductive maturity, they begin allocating energy previously used for growth into reproduction; and 4) juvenile mortality rates keep population density constant, i.e. *R*_*0*_ = 0. In Charnov’s model, selection on age at maturity balances the trade-off between maturing later, thus having more energy to invest in reproduction due to increased size at maturity, versus maturing earlier and accordingly increasing the probability of surviving to reproduction (Charnov 1993). Organisms with high adult mortality rates thus must mature quickly in order to reproduce before dying, while organisms with lower mortality rates can afford to invest more time and energy into growth. However, since life history parameters generally have units of mass and time, this sort of cost-benefit approach to life history theory is ill-equipped to compare small, short-lived organisms to large, long-lived organisms. In order to compare and classify such diverse life history strategies, Charnov proposed using dimensionless ratios and products that eliminate the mass- and time-dependence of life history parameters. Since these metrics lack units, they allow comparisons between organisms that may vary widely on those continua, e.g. from the largest whales to the smallest frogs. Empirical evidence based on life history traits from 64 species of mammals has provided general support for Charnov’s evolutionary predictions (Purvis and Harvey 1995), but they have not been previously tested in a broader macroevolutionary context.

Charnov proposed a classification of life histories is based on three particular dimensionless metrics which represent fundamental trade-offs which organisms must navigate in order to maximize fitness (Charnov 2002): lifetime reproductive effort (LRE), relative reproductive lifespan (RRL), and relative offspring size (ROS). The first of these, LRE (lifetime reproductive effort), can be interpreted as the proportion of adult mass that a female will allocate to offspring over her lifespan (Charnov 2002). LRE represents the relationship between that mass investment and adult mortality (the inverse of lifespan), thus measuring the cost of reproduction, one of the most crucial life history trade-offs (Stearns 1989). The second dimensionless metric, RRL (relative reproductive lifespan), quantifies the trade-off between the time spent preparing to reproduce and the total amount of time available for reproduction (Charnov 1993, 2002). The final of Charnov’s dimensionless metrics, ROS (relative offspring size), measures the relative size at which offspring can occupy the adult niche without parental care (Millar 1977). Charnov hypothesized that variation in these metrics would be greater between major groups of organisms than within groups, since organisms within a group share similar life history constraints (Charnov 2002).

We hypothesize that variation in these life history metrics between major groups of tetrapods has been driven by three key adaptations that impact adult survival and allocation to reproduction – the amniotic egg, endothermy, and flight. The amniotic egg contains membranes improving gas exchange, which allow amniotes to produce larger eggs and maintain much higher rates of egg respiration than amphibians (Thompson and Russell 1998). The increased size of the amniotic egg and the more substantial yolks they contain (Romer 1957) allow for more developed offspring, potentially increasing juvenile survival rates and reproductive allocation in amniotes. Endothermy enables organisms to attain greater metabolic power and potential for production (Gillooly et al. 2002) and exploit a wider range of environments (Rolland et al. 2018), but it is also energetically costly to maintain and limits the minimum size of endotherms. Without these constraints, ectotherms can potentially display a wider variety of life history strategies in response to their local environmental conditions. Finally, the evolution of flight reduces predation risk on volant organisms, decreasing their extrinsic mortality rates and lengthening their lifespans (Holmes and Austad 1994; Healy et al. 2014). Flight also imposes higher parental investment costs (Farmer 2000), and flight itself is energetically costly, which can potentially impact the availability of resources for reproduction. By altering constraints on investment in reproduction and survival, these key adaptations may have impacted the evolution of life history strategies.

Here we use Charnov’s dimensionless life history metrics to assess how these evolutionary innovations altered life history strategies of tetrapods. To conduct comparative analyses between groups of organisms, Charnov envisioned a “life-history cube” defined by LRE, RRL, and ROS as axes, with different groups of organisms occupying different regions of this trait-space (Charnov 2002). Since these three dimensionless metrics are hypothetically uncorrelated with body mass in at least some taxa, we introduced mass as a fourth axis. We employ hypervolume algorithms (Blonder et al. 2017) to create a four-dimensional life history “cube” to test whether major evolutionary transitions in metabolism, physiology, and ecology produce systematic variation in organisms’ life history strategies.

## Methods

### Data

We compiled life history trait data for birds, mammals, reptiles, and amphibians from multiple sources to calculate the dimensionless life history metrics. For the birds and mammals, we used data exclusively from the Amniote Life History Database (Myhrvold et al. 2015). For the reptiles, we supplemented the data available in Amniote with another published set of reptile life history traits (Allen et al. 2017) through a two-step process. First, if a reptile species present in the Amniote database lacked trait data for one of the life history traits necessary to calculate the dimensionless metrics, we filled in the corresponding value from Allen et al. (2017). Secondly, we added trait data for species present in the Allen et al. database but not in Amniote. For the amphibians, we obtained life history trait data from the AmphiBIO database (Oliveira et al. 2017).

### Calculation of Dimensionless Metrics

We used the combined amniote and amphibian data to calculate the three dimensionless life history metrics for 1,650 tetrapod species, including 171 birds, 842 mammals, 491 reptiles, and 113 amphibians.

#### Lifetime Reproductive Effort (LRE)

The first dimensionless metric, LRE, is the product of reproductive effort and average adult lifespan. Charnov defines reproductive effort as *R*/*m*, where *R* is the organism’s average reproductive allocation per unit time and *m* is the average adult body mass (Charnov 2002). To calculate *R*, we multiplied litter or clutch size by the number of litters or clutches per year, yielding the number of offspring per year, and then multiplied this value by the mass of offspring at independence. AmphiBIO reports a minimum and maximum clutch size, so we averaged these two values when calculating *R*. We defined independence as fledging for birds, weaning for mammals, hatching for reptiles, and offspring or egg for amphibians (whichever value for offspring mass was provided in AmphiBIO). After calculating R, we divided by the average adult body mass for the amniotes and the maximum adult body mass for the amphibians to calculate reproductive effort. While Charnov’s model calls for an average body mass, AmphiBIO only provides a maximum adult body mass, so we used this value to provide an approximate value of this metric for amphibians (Oliveira et al. 2017). Finally, we multiply reproductive effort by adult lifespan to calculate LRE. We used maximum longevity, rather than average longevity, for all classes due to data quality and availability.

#### Relative Reproductive Lifespan (RRL)

To calculate RRL, we divided adult lifespan by the time to female maturity. Since Amniote reports longevity in years and age at female maturity in days, we converted longevity to days to keep the ratio dimensionless. AmphiBIO reports minimum and maximum age at sexual maturity, so we averaged these values to calculate an average.

#### Relative Offspring Size (ROS)

In order to calculate the final dimensionless metric, ROS, we divided mass at independence by average adult body mass. We used the same criteria for independence for birds, mammals, reptiles, and amphibians as used to calculate R. Since AmphiBIO reports offspring size as minimum and maximum lengths rather than mass, we used allometry equations for *Anura* and *Caudata* to convert these lengths to offspring mass (Santini et al. 2017). We used the models predicting mass from SVL for *Anura* and *Caudata*, rather than those including habitat and paedomorphy since the majority of frog species in AmphiBIO existed in multiple habitats and paedomorphy was not reported for the salamander species. After converting the minimum and maximum lengths at independence to mass, we averaged these two masses to calculate mass at independence for the amphibians.

### Correlated Evolution Analyses

We first examined whether the dimensionless metrics exhibit correlated evolution with adult body mass. For the mammals, we performed phylogenetic analyses using the Fritz et al. species-level supertree with the best date estimates (Fritz et al. 2009). For the birds, we used the dated phylogeny of extant bird species published by Jetz et al. (2012). We constructed our bird phylogeny using the Hackett et al. (2008) backbone (Jetz et al. 2012). Since reptiles are a paraphyletic group, we restricted our phylogenetic analyses to Squamata, the most diverse reptile order. To analyze this order, we used a time-calibrated phylogeny of squamates (Zheng and Wiens 2016). Finally, for the amphibians, we used a congruified time-tree from the PhyloOrchard package (O’Meara et al. 2013), using the Alfaro et al. timetree of gnathostomes (Alfaro et al. 2009) as the reference and the Pyron and Wiens amphibian phylogeny as the target (Pyron and Wiens 2011).

We stitched together these four phylogenies to create a tetrapod phylogeny to visualize LRE, RRL, and ROS across all four classes. We used divergence estimates from the TimeTree of Life (Hedges et al. 2006) to combine the individual phylogenies according to a pipeline used by Uyeda *et al*. (2017).

We performed phylogenetic least-squares regression analyses (PGLS) (Grafen 1989) using the phylogenies listed above to determine how much of the variation in the relationship between each of the three dimensionless metrics and body mass can be explained by evolutionary relationships. We conducted PGLS using the branch-length transformation indicated by the best-fit model of body mass evolution for each class. We fit Brownian motion, Ornstein-Uhlenbeck, Pagel’s lambda, and kappa models of log body mass using the GEIGER package in R (Harmon et al. 2008). To determine the best-fit model, we ranked by Akaike Information Criterion (AIC) and selected the model with the lowest AIC value.

### Hypervolume Analyses

We used hypervolumes (Blonder et al. 2014, 2017) to investigate the way that different groups of organisms vary with respect to their life history strategies. For the species with complete data coverage, we created four-dimensional hypervolumes with adult body mass and the three dimensionless metrics as axes. All four axes were log-transformed for analysis. We generated hypervolumes for the four major classes of tetrapods: birds, mammals, reptiles, and amphibians. All hypervolumes were created using the hypervolume R package using the Gaussian KDE method with the default Silverman bandwidth estimator (Blonder et al. 2017). To compare hypervolumes between groups, we calculated individual volumes and pairwise overlap metrics, including Sorensen similarity and the fraction of unique volume. The size of these hypervolumes represents the total diversity of life history strategies for a given class, while the overlap indicates the similarity in strategies between classes.

Code for all analyses, as well as data files, can be found at https://github.com/KerkhoffLab/bodymasspatterns (see Supplementary Information and Figures).

## Results

### Metric values across classes

We compared values for the three dimensionless metrics across the four tetrapod classes to examine variation in life history strategies across the major groups. LRE showed a distinct increase across the classes, from amphibians to reptiles to mammals, and finally to birds (ANOVA; F=1017.8; d.f.=3; p<2.2×10^−16^) (Fig. 1B). ROS demonstrated a similar pattern to LRE: mean log ROS differed across all four classes (Tukey HSD; p<0.05), increasing by 98.1% from amphibians to birds (Fig. 1D). RRL also differed between all four classes (ANOVA; F=242.9; d.f.=3; p<2.2×10^−16^) (Fig. 1C). Mammals had the highest mean RRL value, followed by birds, reptiles, and amphibians (Tukey HSD; p<0.05).

**Figure 1.**
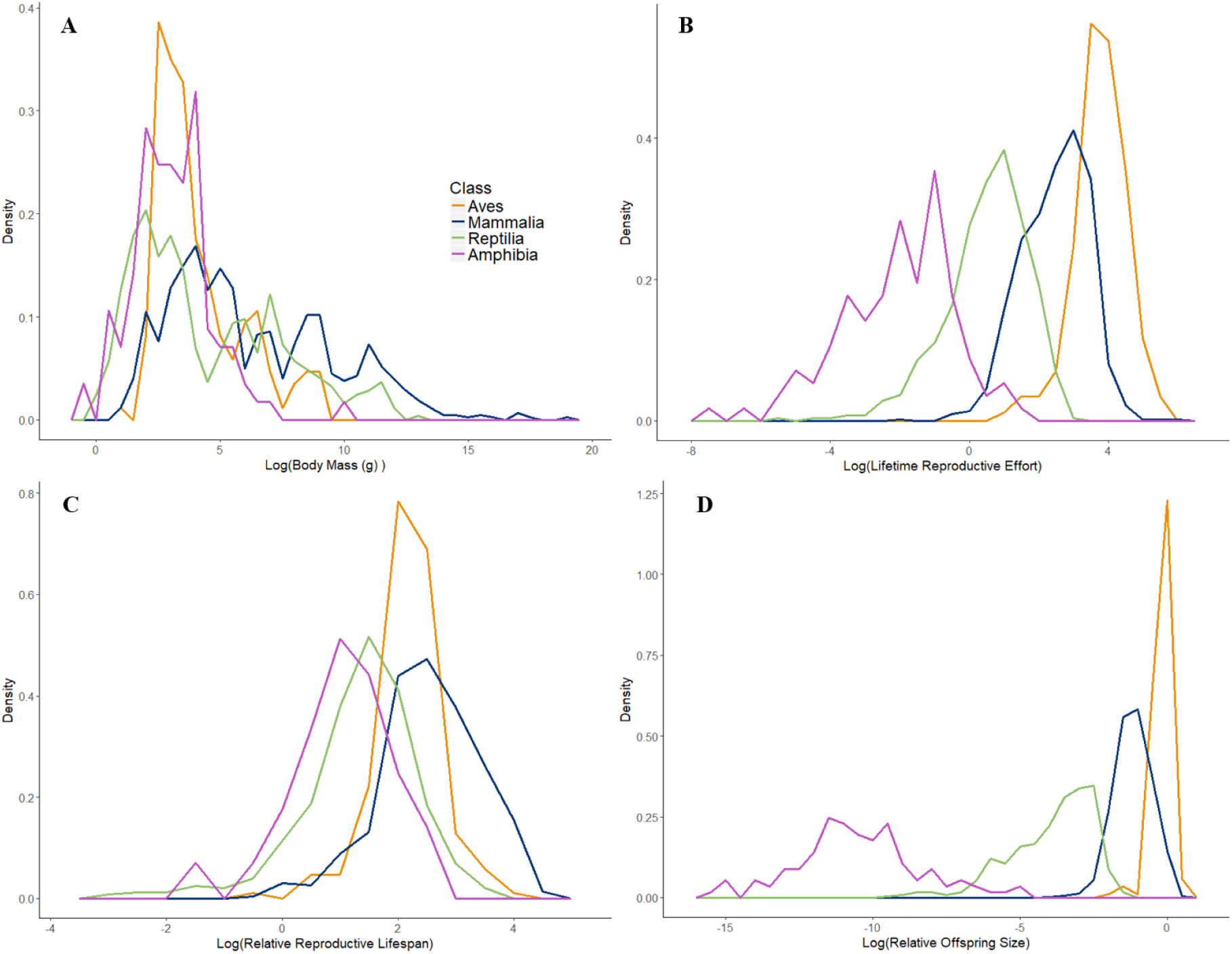
Frequency polygons of (A) log-transformed body mass, (B) LRE, (C) RRL, and (D) ROS for tetrapod species with values for all three of the dimensionless traits.When viewed on the tetrapod phylogeny, LRE shows distinct shifts in values across the four classes: higher values of LRE appear to have evolved both in birds and mammals, while the amphibians display consistently low values (Fig. 2A). In comparison, RRL does not show as clear distinctions between classes, but, in general, the lowest values are found in amphibians and reptiles (Fig. 2B). There are certain clades of mammals, however, like the family Soricidae, which have low RRL values comparable to or lower than those found in amphibians and squamates. In the birds, as well, certain Charadriiformes have quite low RRL values. Of the three metrics, ROS shows the most dramatic shifts across the four classes of tetrapods (Fig. 2C), with the lowest values in amphibians, followed by squamates, and finally mammals and birds.

### Correlated Evolution

Next, we performed phylogenetic analyses to investigate possible evolutionary mechanisms that may influence the relationship between these metrics and body mass across different clades. Based on the AIC values, we used the Ornstein-Uhlenbeck model for the ectotherms, and Pagel’s lambda model for the endotherms. After accounting for evolutionary relationships, body mass was negatively correlated with LRE in amphibians and mammals (p<0.05), but there was no significant relationship for the squamates and birds (p>0.05) (Table 1). RRL, on the other hand, was only significantly correlated with body mass in the squamates (p=0.0155) (Table 1). Body mass was negatively correlated with ROS across all the clades after accounting for phylogeny (p<0.05). The magnitude of the slope of this relationship decreased from the oldest class (Amphibia) to the youngest (Aves) (Table 1).

**Table 1.**
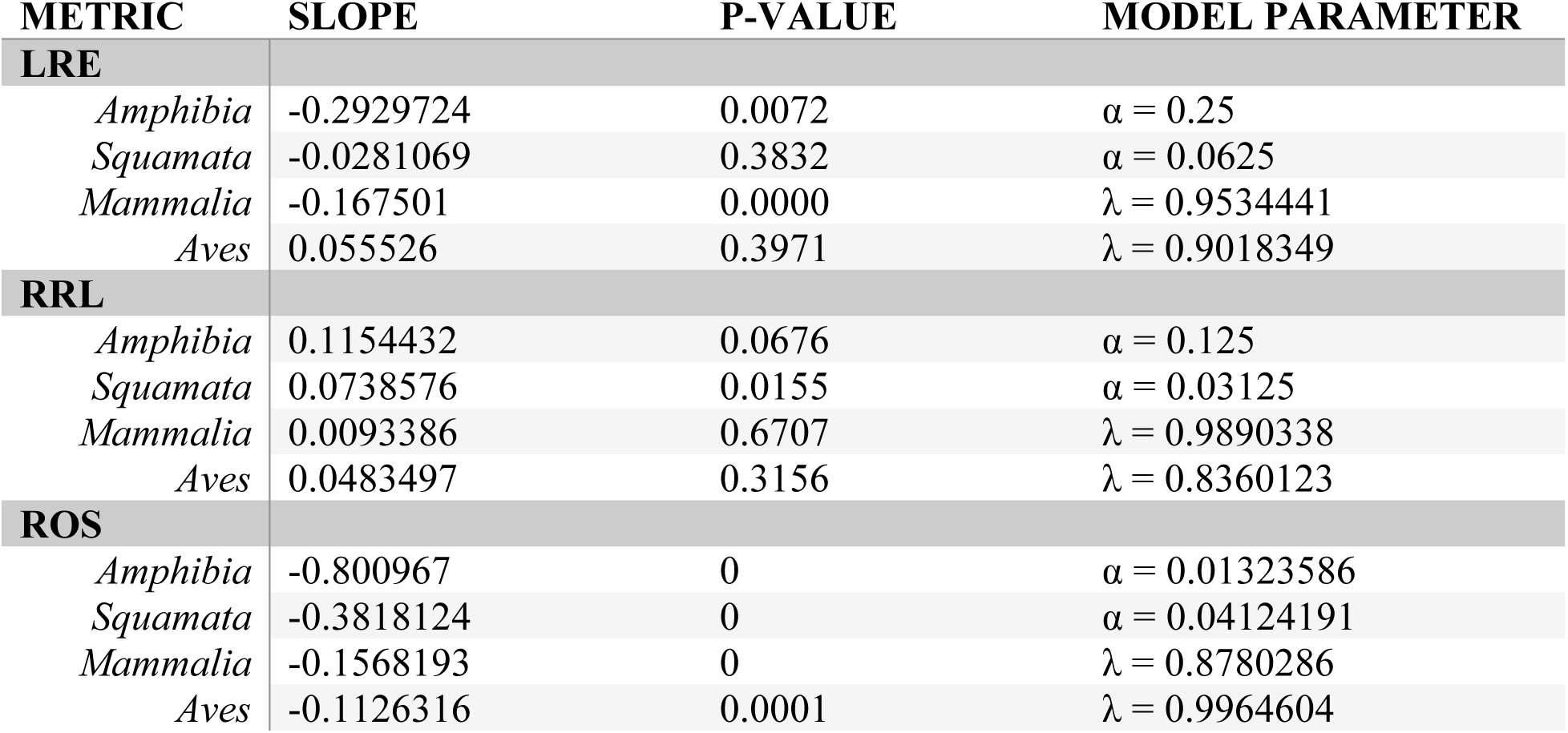
Coefficients for phylogenetic least-squares analysis (PGLS) between each log-transformed dimensionless metric and log body mass for species with trait values for all three metrics and body mass. PGLS was performed using a correlation matrix based on the model of body mass evolution with the lowest AIC for each class: Ornstein-Uhlenbeck for amphibians and squamates and Pagel’s lambda for mammals and birds.

| METRIC | SLOPE | P-VALUE | MODEL PARAMETER |
| --- | --- | --- | --- |
| **LRE** |  |  |  |
| *Amphibia* | -0.2929724 | 0.0072 | $\alpha = 0.25$ |
| *Squamata* | -0.0281069 | 0.3832 | $\alpha = 0.0625$ |
| *Mammalia* | -0.167501 | 0.0000 | $\lambda = 0.9534441$ |
| *Aves* | 0.055526 | 0.3971 | $\lambda = 0.9018349$ |
| **RRL** |  |  |  |
| *Amphibia* | 0.1154432 | 0.0676 | $\alpha = 0.125$ |
| *Squamata* | 0.0738576 | 0.0155 | $\alpha = 0.03125$ |
| *Mammalia* | 0.0093386 | 0.6707 | $\lambda = 0.9890338$ |
| *Aves* | 0.0483497 | 0.3156 | $\lambda = 0.8360123$ |
| **ROS** |  |  |  |
| *Amphibia* | -0.800967 | 0 | $\alpha = 0.01323586$ |
| *Squamata* | -0.3818124 | 0 | $\alpha = 0.04124191$ |
| *Mammalia* | -0.1568193 | 0 | $\lambda = 0.8780286$ |
| *Aves* | -0.1126316 | 0.0001 | $\lambda = 0.9964604$ |

### Hypervolumes

Even though the four tetrapod classes overlap in several of the individual life history metrics as well as body mass, they occupy distinct regions of multidimensional life history space. As predicted by Charnov’s life history “cube” (Charnov 2002), the four-dimensional hypervolumes for each class vary in size, shape, and area of trait space occupied (Fig. 3). The volume of each hypervolume, representing diversity in trait combinations present for a given group, decreased dramatically across classes in order of evolution. The two ectothermic hypervolumes, Amphibia and Reptilia, were 4.81 and 3.06 times larger, respectively, than the mammal hypervolume, while the bird hypervolume was 17% of the size of the mammal hypervolume. The class hypervolumes also occupied highly distinct regions of the trait space. The birds and mammals had the most overlap, still with a Sorensen similarity of only 0.1392 (Table 2). The amphibian hypervolume was the most unique, not overlapping with the endotherm hypervolumes at all, and only having a Sorensen similarity of 0.0271 with the reptiles (Table 2).

**Table 2.**
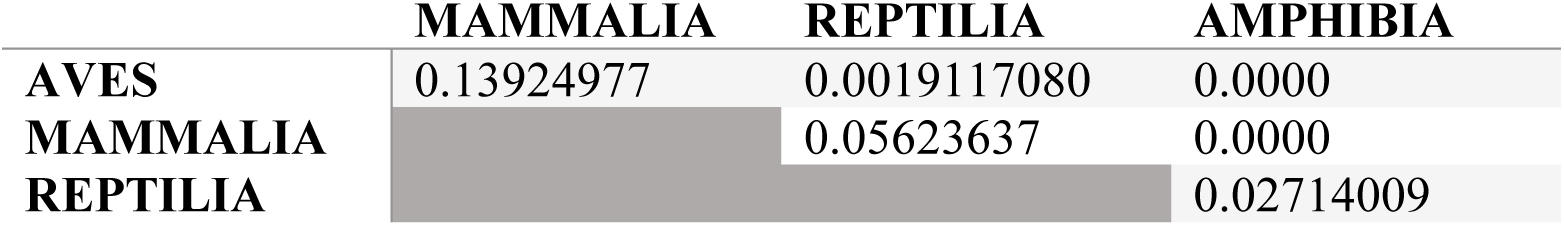
Sorensen similarity of hypervolumes between classes.

**Figure 2.**
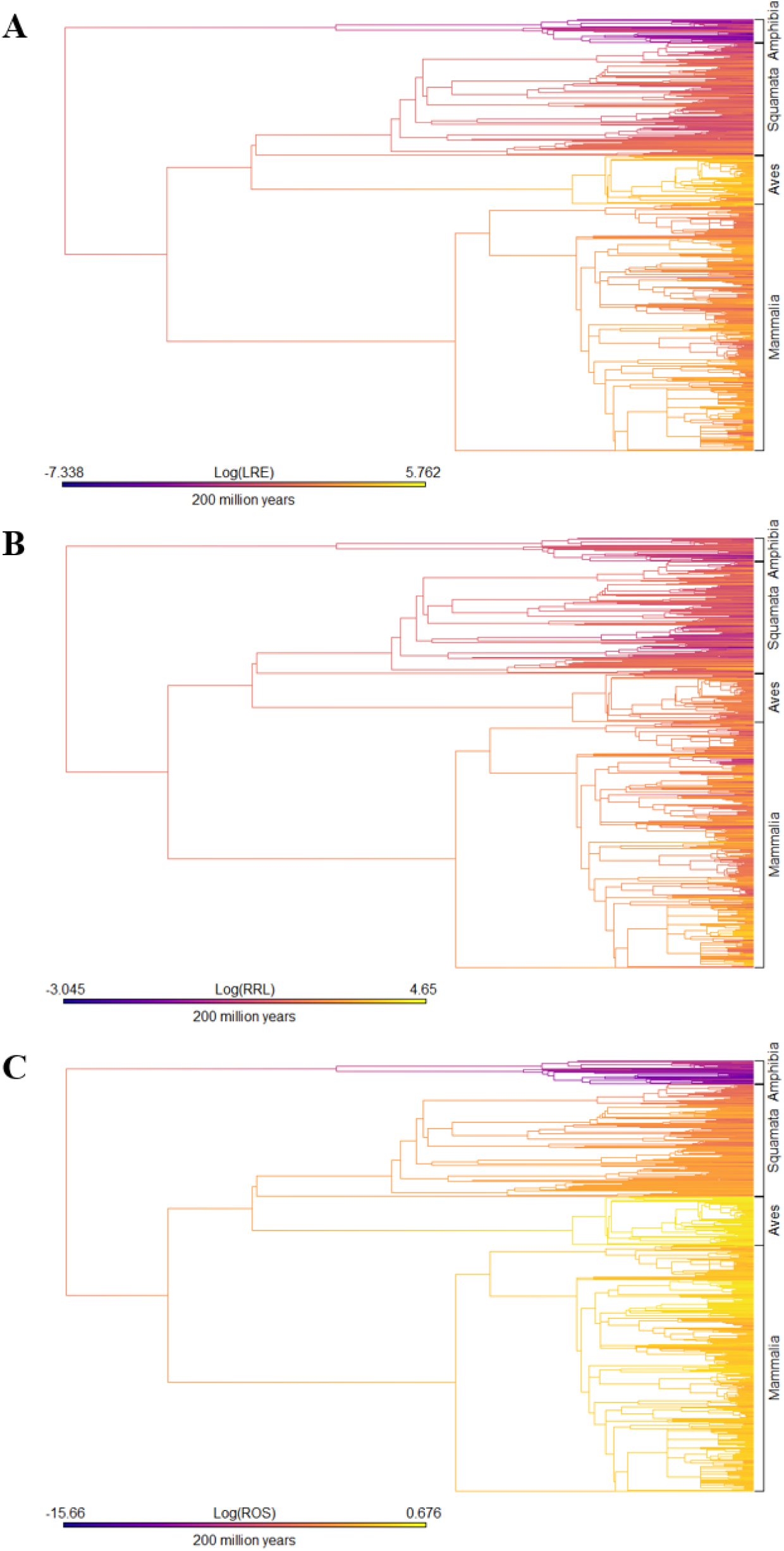
Plots of (A) LRE, (B) RRL, and (C) ROS across a supertree of tetrapods. Trait values along the edges and at nodes were estimated based on a Brownian motion model of evolution. The color ramp bar serves as a legend for trait values and a scale for branch lengths.

**Figure 3.**
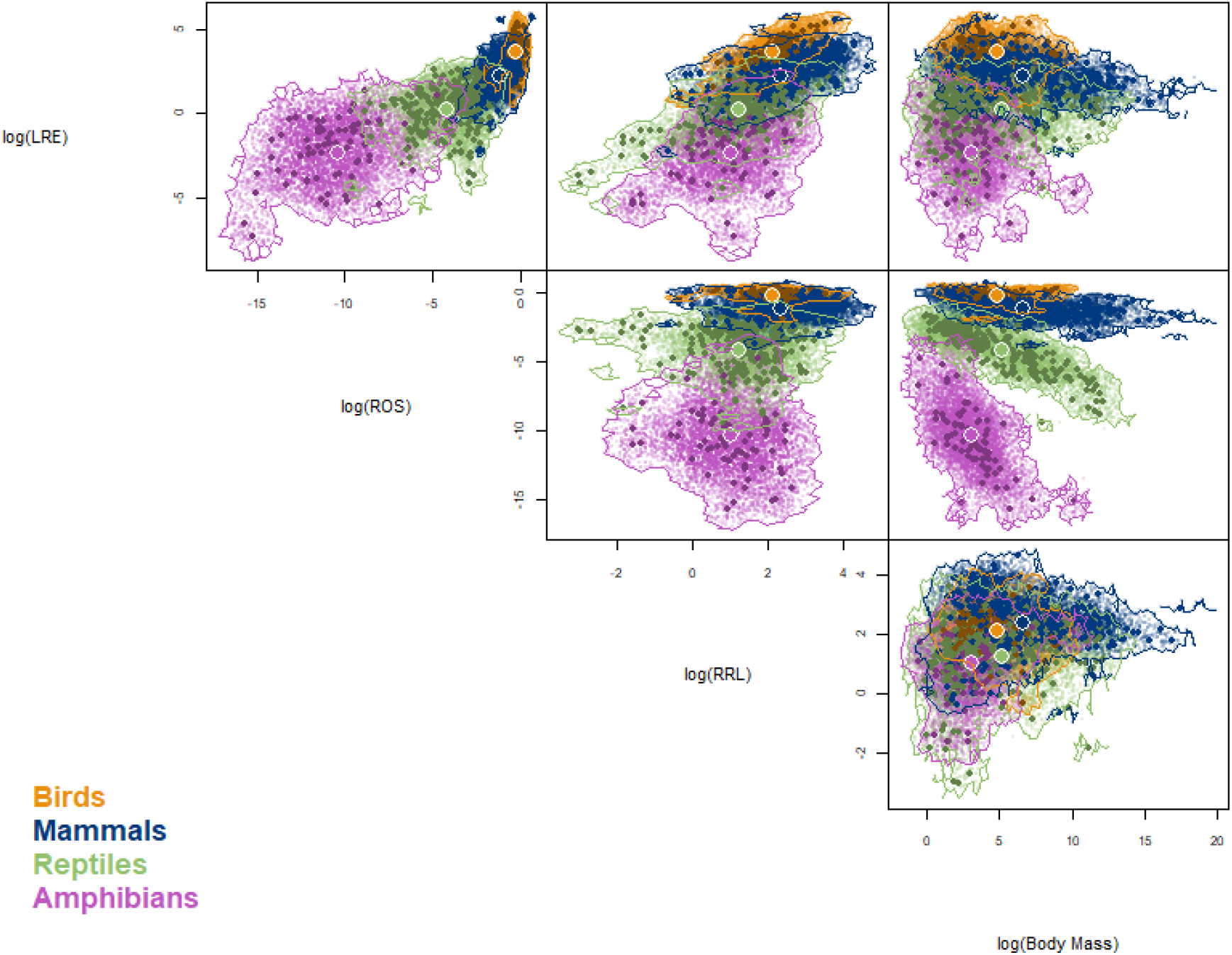
Gaussian hypervolumes for the four classes of tetrapods. Large colored points represent the centroids of each hypervolume. Small dark points represent trait values for individual species while small light points represent random points. Volume of bird hypervolume is 36.69, mammal is 214.98, reptile is 657.93, and amphibian is 1034.54.

## Discussion

Charnov’s dimensionless life history metrics, (LRE, RRL, and ROS) provide a novel framework to compare organisms’ life history strategies. The three dimensionless metrics show a range of patterns of correlated evolution, which drive their relationships with body mass in extant species. Furthermore, the major tetrapod classes display unique combinations of these metrics (Figure 3). The differences in subspaces occupied by each class may reflect the effects of crucial evolutionary transitions in energetics, physiology, and ecology. Thus, LRE, RRL, and ROS enable us to compare life history strategies. By observing how the metrics change in tandem with major adaptations, such as evolution of the amniotic egg, endothermy, and flight, we explore the possible ecological and evolutionary influences on life history.

### Amphibians to amniotes: evolution of the amnion

Amphibians are notably different from the amniote groups (reptiles, birds, and mammals) in their life histories. Amphibians take longer to reach reproductive maturity relative to their total reproductive lifespan (lowest mean RRL; fig 1c), invest much less in reproduction (lowest mean LRE; fig 1b), and produce smaller offspring at independence relative to adult size (lowest ROS; fig 1d). Many of these differences reflect the constraints imposed by a non-amniote egg. Amniotic eggs contain unique membranes which allow them to be much larger and maintain higher rates of respiration than those of amphibians, which are limited by the rate of diffusion of oxygen through the egg (Seymour and Bradford 1995; Thompson and Russell 1998). Because offspring can spend longer in the egg, amniotes supply their eggs with substantial yolks that allow offspring to develop to a greater degree before hatching (Romer 1957). These adaptations allow amniotes to emerge at a higher stage of development than amphibian offspring do (Romer 1957) and may help amniotes reach reproductive maturity more quickly—and spend a greater proportion of their total lifespan reproducing—compared to amphibians. These impacts of the amniote egg on the size of the offspring and the extra investment parents provide their offspring in yolk may also explain why amniotes exhibit higher levels of lifetime reproductive investment and relative offspring size than amphibians (Fig 1B, D).

While the vast majority of comparative life history research focuses on amniotes (Western and Ssemakula 1982; Shine 2005; Sibly et al. 2012; Healy et al. 2014; Capellini et al. 2015), our work highlights the incredible diversity of amphibian life history strategies compared to other tetrapods. Amphibians occupy a region of the life history trait space that is almost completely unique from the other three classes. Moreover, the amphibian life history hypervolume was the largest in volume by almost an order of magnitude (Fig 3), indicating that amphibians possess remarkable diversity in their strategies, especially considering that the hypervolume axes are all log-transformed. This diversity may be driven by the variety of modifications to amphibian life cycles, from neotony to viviparity, which permit variation in clutch size and offspring size (Wilbur and Collins 1973).

### Endotherms vs. ectotherms: the impact of thermic strategy

While the amphibians exhibit the greatest range of life history metric combinations, both of the ectothermic classes occupy regions of trait space many orders of magnitude larger than those occupied by the endothermic classes (Fig 3). This pattern is consistent with the hypothesis that while endothermy conveys advantages (Rolland et al. 2018), it also comes with costs and constraints that can limit life history strategies (Tinkle et al. 1970; Allen et al. 1999; Shine 2005). Endothermy is related to higher metabolic power (Uyeda et al. 2017) and potential for production and an enhanced ability to maintain activity under a broader range of conditions (Crompton et al. 1978; Rolland et al. 2018). These advantages may allow endotherms to have more resources for reproduction (Farmer et al. 2003) or decrease adult mortality through impacts on foraging durations and predator avoidance (Bennett and Ruben 1979; Bennett 1991). However, endothermy is energetically expensive (Bennett and Ruben 1979; Koteja 2004) and it is especially difficult for small organisms to maintain the thermal differential with their environment that endothermy requires. These constraints and advantages have the potential to alter the viability of different life history strategies via their impact on reproductive allocation and survival. Our results suggest that these advantages and constraints conferred by endothermy have resulted lower flexibility and variability among endothermic species in their life history strategies.

The life history pattern in relative offspring size (ROS) exhibits the strongest constraint in endotherms. Mammals and birds produce offspring that must attain a greater proportion of their adult mass before independence, which requires greater parental investment. Not only is mean ROS higher for endotherms than ectotherms, the endotherms also demonstrate much less variation, implying that endothermy may constrain the possible range of ROS. The need for greater offspring size for endotherms could reflect that the fact that thermogenic tissue is expensive to produce and that offspring may need greater levels of parental investment to produce it (Case 1978). It is also difficult–and energetically costly–for small individuals to maintain a thermal differential with the environment which could necessitate greater parental investment to help offspring reach a size that reduces their thermoregulatory costs (Farmer et al. 2003; Shine 2005). Because resources for reproduction are limited, increases in offspring investment generate decreases in the number of offspring that can be produced (Smith and Fretwell 1974). Thus, this need to invest in larger offspring may preclude endotherms from life history strategies that produce many small, mostly independent offspring with minimal parental care. Without the constraints imposed by endothermy, ectotherms can take on a wider range of life history traits, including decreased offspring size and increased fecundity (Tinkle et al. 1970; Allen et al. 1999; Shine 2005).

While we found an increased diversity of life history strategies in ectotherms, other studies have shown that amphibians and reptiles have a slower rate of environmental niche evolution than birds and mammals (Rolland et al. 2018). Previous research has highlighted a similar relationship between life history flexibility and environmental niche. An increase in the number of annual life history stages in adults (e.g. breeding, migration, molting, etc.) allows the organism to maximize fitness across a greater range of environmental conditions (Wingfield 2008). Mammals and especially birds exhibit more of these stages than ectotherm clades. Having a large number of these stages, however, limits life history flexibility in terms of timing (Wingfield 2008). We find a similar trade-off: while endotherms can tolerate a much wider range of climates (Rolland et al. 2018), ectotherms exhibit much more flexibility in life history strategies. These results suggest that, while endothermy offers organisms options in terms of environmental niches (Rolland et al. 2018), it also imposes certain constraints on life history.

### Taking flight: the importance of volancy

Volancy drives changes in longevity and parental investment that impose strong constraints on the three dimensionless metrics. Lifespan is longer in birds, as well as in volant mammals, compared to non-volant mammals (Holmes and Austad 1994; Healy et al. 2014). Research has suggested that flight enables organisms to escape predation more successfully (Pomeroy 1990; Holmes and Austad 1994), which in turn decreases extrinsic mortality and increases longevity. Volancy also affects life history via its impact on parental care. In general, volant species must allocate more energy to parental care in order to supply young with food prior to independence (Farmer 2000). These effects of flight lead to a variety of changes in the three metrics in both birds and bats, although these two clades also face separate constraints that affect their life history strategies differently.

Despite the increase in longevity driven by flight, birds have slightly lower RRL values than all mammals, including bats. If birds had similar ages at female maturity as mammals, we would expect RRL to be higher in birds due to this longer lifespan. We observed the opposite, however, with mammals having a significantly higher mean RRL value, indicating that birds take relatively longer to mature despite having longer overall lifespans. Volancy itself does not appear to cause lower RRL, however, since bats have RRL values more similar to mammals than to birds. Instead, it appears as though birds have unique constraints on breeding not faced by mammals. Almost all small birds wait at least one year from hatching before they begin breeding, while small mammals do not. Several factors may play a role in this discrepancy. First, migration is more common in small birds than in small mammals, and this additional life history stage constrains the timing of breeding (Wingfield 2008). Second, even in non-migratory species, birds have a higher metabolic rate and body temperature than mammals (Nagy 1987). Since reproduction typically necessitates an additional increase in metabolic rate (Farmer 2000), birds have more seasonal constraints on breeding due to the need to maintain the incredibly high energy investment in breeding. Finally, since birds have lower mortality rates in general, they may be able to afford a longer period of investment in growth before beginning reproduction, as predicted by Charnov’s evolutionary model (Charnov 1993). This collection of factors may contribute to lower avian RRL values compared to mammals.

Birds display the highest LRE values of all four tetrapod classes. Volancy is the most important driver of increased lifespan in endotherms, which leads to increases in LRE by increasing the amount of time birds can devote to reproduction over their lifetime (Healy et al. 2014). Furthermore, birds have the highest metabolic rates and body temperatures of all tetrapods (Nagy 1987). Reproduction necessitates a further metabolic increase beyond this already high investment (Farmer 2000). The elevated LRE of birds may reflect that extra cost of avian reproduction compared to that of mammals and ectotherms.

Flight appears to be a strong constraint on mean relative offspring size (ROS) values. Bats occupy a range of ROS values much more similar to that of birds than other mammals (Fig. S-1). In terms of the trait space defined by the three dimensionless metrics, they resemble birds far more than mammals on the ROS axis, while behaving much more like mammals in their range of LRE values (Fig. S-2). This evidence suggests that flight operates particularly on ROS out of the three metrics: volant organisms must approach adult body mass before they can be independent from their parents.

### Relationship between lifetime reproductive effort and relative offspring size

As both lifetime reproductive effort (LRE) and relative offspring size (ROS) increase across the four tetrapod classes from amphibians to birds, these two metrics demonstrate a striking positive correlation (Fig. 3). Tetrapods with high reproductive effort and high longevity tend to have higher masses at independence relative to their adult body mass and vice versa. As clades have evolved successive adaptations, from the amniotic egg to endothermy, and finally volancy, organisms have moved towards these higher reproductive investments and relative masses at maturity. This relationship is constrained in two ways. First, ROS is a component of LRE, which calculates reproductive effort as the product of ROS, longevity, and the number of offspring per year. Thus the slope of the correlation between LRE and ROS is approximately the average number of offspring per year across an organism’s reproductive lifespan. This value is proportional to average reproductive output R0, i.e. fitness. R0 is hypothesized to be the product of survival to adulthood and this value, which is the ratio of LRE to ROS (Charnov et al. 2007).

In the log-log plot of LRE vs. ROS, we observe higher intercepts in the endotherm classes compared to the ectotherm classes, indicating that the endotherms may have higher R_0_ values. However, we also observe increasing slopes of the log-log plot moving from amphibians through birds, indicating a more complex interaction between these two metrics. Additionally, the case study of bats indicates that other factors influence this relationship. Bats resemble birds in their constraints on ROS, but they exhibit LRE values like those of other mammals (Fig. S-2). Further study of the relationship between LRE and ROS may provide a method to compare relative fitness between and within classes.

### Conclusions

The Charnov dimensionless life history metrics provide a useful tool for exploring the causes and consequences of life history variation among clades. These metrics have, in most cases, evolved with body mass in a correlated fashion within classes. Nonetheless, lifetime reproductive effort (LRE), relative reproductive lifespan (RRL), and relative offspring size (ROS) provide a highly informative lens through which to investigate life history strategies across a wide range of organisms. The four tetrapod classes differ drastically in their combinations of LRE, RRL, and ROS, indicating that they adopt different strategies to address fundamental life history trade-offs. We find that ectotherms have tremendous variation in life history strategies, while birds are highly constrained relative to even mammals. These distinctions imply that major adaptations in energetics, physiology, and ecology result in changes to the dimensionless metrics, and, by extension, the underlying trade-offs they represent.

## Acknowledgments

We are very grateful to Dr. Natalie Wright for her invaluable advice on interpreting PGLS, as well as comments on the manuscript. Additionally, many thanks to Dr. Benjamin Blonder for assistance with the hypervolume figure. Also, Dr. Elizabeth Schultz provided very helpful feedback on avian life history trends specifically. Finally, thank you to Dr. Benjamin Blonder, Dr. Brian Maitner, and Dr. Brian Enquist for feedback on the early stages of the project and to Dr. Eric Charnov for encouragement in exploring dimensionless life histories.

This research was funded by National Science Foundation (NSF) award DEB-1556651 and the 2017 Kenyon Summer Science Scholarship.

## Supplementary Information and Figures

### Data and Code

R code for all figures and analyses, as well as data and links to data files can be found at https://github.com/KerkhoffLab/bodymasspatterns.

**Figure S-1.**
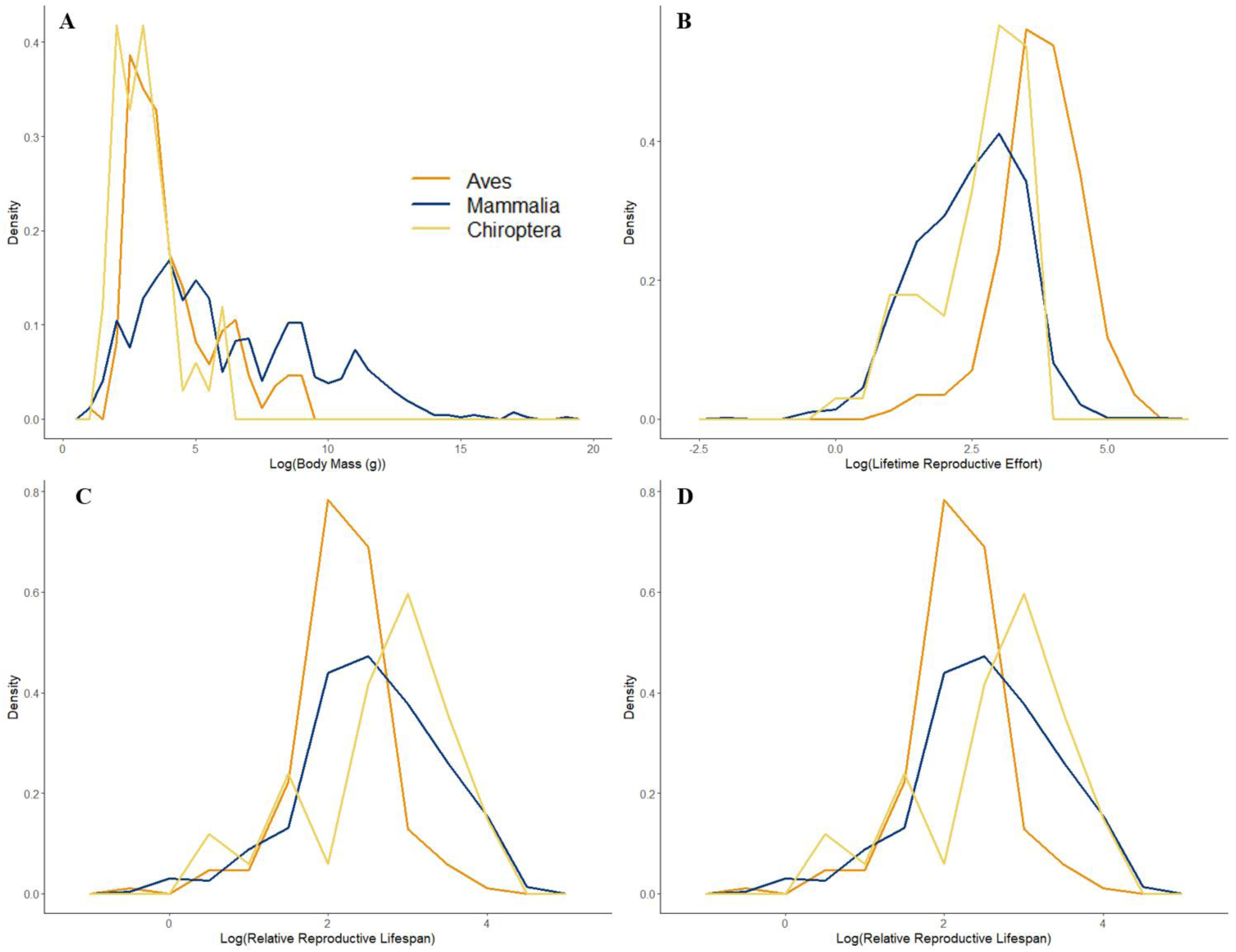
Frequency polygons of (A) log-transformed body mass, (B) LRE, (C) RRL, and (D) ROS for bird, mammal, and bat species with values for all three of the dimensionless traits.

**Figure S-2.**
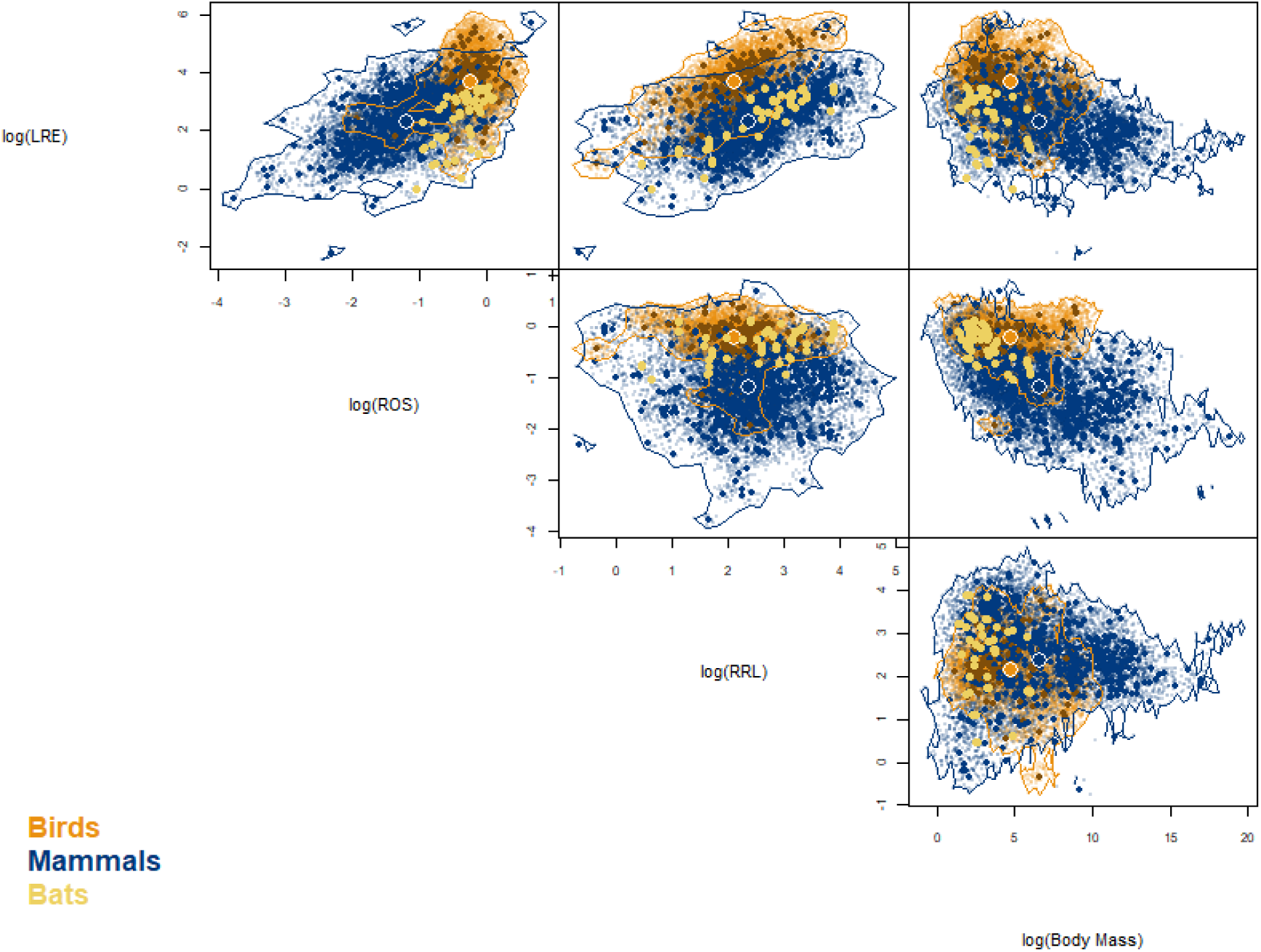
Bat metric values overlaid on Gaussian hypervolumes of birds and mammals. Large colored points represent the centroids of each hypervolume. Small dark points represent trait values for individual species while small light points represent random points.

**Table S-1.** Sorensen similarity of hypervolumes between birds, mammals, and bats.

|  | AVES | MAMMALIA |
| --- | --- | --- |
| CHIROPTERA | 0.1924 | 0.2807 |
| AVES |  | 0.1392 |

